# Vertical transmission of *Wolbachia* bypassing the germ line in an aphid

**DOI:** 10.1101/2025.04.14.648810

**Authors:** Tomonari Nozaki, Yuuki Kobayashi, Shuji Shigenobu

**Affiliations:** Laboratory of Evolutionary Genomics, National Institute for Basic Biology (NIBB), Okazaki, Aichi 444-8585, Japan; Department of Basic Biology, School of Life Science, The Graduate University for Advanced Studies, SOKENDAI, Okazaki, Aichi 444-8585, Japan

**Keywords:** Maternal transmission, Endosymbiotic bacteria, Viviparous reproduction, *Wolbachia*, Cedar bark aphid

## Abstract

*Wolbachia*, a widespread endosymbiotic bacterium that infects a broad range of arthropods and nematodes, relies on vertical transmission from mother to offspring. This process often involves colonization of the host germline, subsequent transfer to developing oocytes, and utilization of host yolk protein transport mechanisms such as vitellogenin uptake. However, the transmission strategies employed by *Wolbachia* in viviparous insects such as aphids are poorly understood. Here, we demonstrate a novel *Wolbachia* transmission mode in the cedar bark aphid *Cinara cedri* that bypasses germline cells. After confirming stable *Wolbachia* infection in *C. cedri*, we visualized the localization of *Wolbachia*, along with the obligate symbionts *Buchnera aphidicola* and *Serratia symbiotica*. Consistent with previous reports, *Wolbachia* in *C. cedri* were predominantly observed within maternal and embryonic bacteriocytes, the specialized cells housing obligate symbionts. Notably, *Wolbachia* cells were rarely detected in germline cells or early-stage embryos and were directly transmitted from maternal bacteriocytes to developing embryos, coinciding with obligate symbiont transfer. These results suggest that *Wolbachia* in *C. cedri* has evolved a unique “piggybacking” strategy, utilizing the obligate symbiont transmission system. Our study highlights the diversity of endosymbiont maternal transmission strategies and provides new insights into the underlying molecular mechanisms of action.

## Introduction

*Wolbachia*, an alphaproteobacterial genus with widespread intracellular symbionts, infects approximately half of all arthropod and nematode species worldwide (Kaur et al., 2021). Its primary mode of transmission is vertical, from mother to offspring (Pietri et al., 2016). Maternal transmission, coupled with its remarkable ability to manipulate host reproduction, including parthenogenesis induction, cytoplasmic incompatibility, male killing, and feminization, is crucial for its evolutionary success (Kaur et al., 2021; Werren et al., 2008). Vertical transmission facilitates rapid dissemination within host populations, ensuring the long-term persistence of *Wolbachia* (Russell et al., 2019). Although detectable in various host tissues, *Wolbachia* typically resides predominantly in the female germline (Kaur et al., 2021; Pietri et al., 2016). Although present in male germlines during early spermatogenesis, *Wolbachia* are subsequently eliminated from sperm cysts (Clark et al., 2002), effectively precluding paternal transmission. Maternal transmission is well-established, occurring through *Wolbachia* accumulation within female germline stem cells and their subsequent transfer from nurse cells to developing oocytes during oogenesis (Guo et al., 2018; Hosokawa et al., 2010; Kaur et al., 2021; Russell et al., 2019).

The mechanisms of maternal transmission have been extensively studied in oviparous insects such as *Drosophila* and mosquitoes (Kaur et al., 2021; Russell et al., 2019). However, the transmission strategies employed by *Wolbachia* in viviparous insects are poorly understood. Viviparous reproduction, an alternative reproductive mode frequently observed among predominantly oviparous insects (Wheeler, 2003), may present unique challenges to *Wolbachia* transmission. For instance, in the viviparous reproduction of aphids, the connection between germline cells and oocytes is limited to a brief and early developmental window, and nutrient supply from nurse cells is reasonably assumed to be restricted (Bickel et al., 2013; Miura et al., 2001). These factors may hinder the well-known *Wolbachia* infection mechanisms. Therefore, we hypothesized that *Wolbachia* in viviparous aphids may employ one or more of the following strategies: (1) highly efficient early germline infection with low bacterial titers, (2) exploitation of alternative nutrient uptake pathways, or (3) vertical transmission through other means, such as direct transmission into the developing embryo. Detailed observations of the *Wolbachia* infection process during viviparous reproduction of aphids will expand our understanding of the diverse mechanisms of symbiont vertical transmission.

Aphids (Hemiptera: Aphididae) are prime model systems for studying intracellular symbiosis (Moran et al., 2008; Shigenobu and Yorimoto, 2022). Almost all aphids rely on *Buchnera aphidicola* (hereafter *Buchnera*), an obligate symbiotic bacterium harbored within hyperpolyploid cells called “bacteriocytes,” for essential nutrients (Nozaki and Shigenobu, 2022). They frequently harbor other obligate or facultative symbiotic bacteria, such as *Serratia symbiotica* (hereafter *Serratia*) (Manzano-Marín et al., 2023; Monnin et al., 2020; Moran et al., 2005) in the bacteriome cells. The phylogenomic status, transmission modes, and functional roles of these symbionts have been extensively investigated (Koga et al., 2012; Oliver et al., 2010; Renoz, 2024). However, the relationship between aphids and *Wolbachia* remains poorly understood largely because of the rarity of stable *Wolbachia* infections in aphids (Augustinos et al., 2011). *Wolbachia* infection has been reported in the banana aphids *Pentalonia nigronervosa* and *Pentalonia caladii* (De Clerck et al., 2014; Jones et al., 2011), and while its functional role in these species is debatable (De Clerck et al., 2015; Manzano-Marín, 2020), a recent study has shown its role in resistance to fungal parasitoids (Higashi et al., 2024). Given that *Wolbachia* strains in the M and N supergroups are primarily associated with aphids (Moreira et al., 2019; Romanov et al., 2020), a unique association between aphids and *Wolbachia* is likely. Therefore, characterizing *Wolbachia* in aphids could provide valuable insights into the evolution of this highly successful insect symbiont.

*Cinara cedri*, the cedar bark aphid, is another species in which *Wolbachia* infections have been widely observed (Augustinos et al., 2011; Gómez-Valero et al., 2004; Jousselin et al., 2016; Nozaki et al., 2022). This aphid is a well-established model of dual-obligate symbiosis that relies on *Buchnera* and *Serratia* for nutritional complementation (Lamelas et al., 2011; Pérez-Brocal et al., 2006). Previous studies have shown that these symbionts, including *Wolbachia*: *Buchnera* and *Serratia* reside in distinct bacteriocytes, and *Wolbachia* is also found within these cells rather than in other tissues or the hemolymph (Gómez-Valero et al., 2004; Manzano-Marín et al., 2016). However, the mode of *Wolbachia* transmission in *C. cedri* remains unclear. In the present study, we characterized the localization and transmission of *C. cedri* symbionts, including *Wolbachia*. We first confirmed consistent *Wolbachia* infection and quantified symbiont abundance using 16S bacterial rRNA amplicon sequencing in Japanese populations. We investigated the localization of the three symbionts (*Buchnera, Serratia*, and *Wolbachia*) within the bacteriome at different embryonic stages. Finally, based on our observations, we propose a novel *Wolbachia* transmission pathway, indicating a “piggyback” mechanism via those of obligate symbionts.

## Materials and methods

### Aphid collection

We collected *C. cedri* from three localities in Japan between 2021 and 2024 (Table 1). When aphids were observed on the twigs of *Cedrus deodara*, they were carefully collected using aspirators or forceps. Almost all individuals collected were viviparous in all seasons, suggesting that they were anholocyclic in Japan (but see SI text). Aphid samples were immediately preserved in 99.5% ethanol for genetic analysis after collection, or dissected and fixed with 4% paraformaldehyde (PFA) for microscopic observation. *C. cedri* has been introduced into several countries, including Japan, along with its host tree, *C. deodara*, which is grown as an ornamental plant (Blackman and Eastop, 2018; Nozaki et al., 2022). We previously confirmed that Japanese colonies of *C. cedri* are genetically homogeneous based on Sanger sequencing of the cytochrome oxidase I regions (Nozaki et al., 2022).

**Table 1.** Sample information of *Cinara cedri* collected in the study.

| # | Locality | Latitude | Longitude | Sample collection | Usage |
| --- | --- | --- | --- | --- | --- |
| 1 | Okazaki, Aichi<br>(NIBB campus) | 34.94769 | 137.16591 | May, 2021 | Amplicon seq. |
|  |  |  |  | Jan., May 2022 | Amplicon seq., Imaging |
|  |  |  |  | Apr., Aug., Oct., Nov. 2023 | Imaging |
|  |  |  |  | May. 2024 | Imaging |
| 2 | Mibu, Tochigi | 36.45498 | 139.80784 | Aug. 2022 | Amplicon seq. |
| 3 | Nishinomiya, Hyogo | 34.72121 | 135.36598 | Aug. 2021 | Amplicon seq. |

### 16S ribosomal DNA amplicon sequencing analysis on *C. cedri* microbiome

To characterize the diversity and relative abundance of bacterial endosymbionts in *C. cedri*, we used 16 individuals from three geographically distinct populations across Japan (six from Aichi, six from Tochigi, and four from Hyogo; Table 1) for high-throughput 16S rRNA sequencing targeting the hypervariable V3/V4 region of the bacterial 16S rRNA gene. Total DNA was extracted as follows: single individuals preserved in 99.5% ethanol were air-dried and quickly rinsed with buffer A (10 mM Tris pH 8.0, 1 mM EDTA, and 25 mM NaCl). Samples in 100 μL buffer A added to 1 μL proteinase K (400 μg/mL) were completely homogenized using BioMasher II (Nippi, Japan). After incubation at 37 °C for 1 h, the samples were heated at 98 °C for 2 min. The samples were preserved at -20 °C until use. Using these total DNA samples, the libraries were constructed according to the 16S rRNA Sequencing Guide “16S Metagenomic Sequencing Library Preparation (15044223 B JPN),” which was provided by Illumina (CA, USA). The V3/V4 region (ca. 460 bp) of the bacterial 16S rRNA gene was amplified using the 16S AmpF_IL and 16S AmpR_IL primers. A 10 μL PCR cocktail contained 2 μL of the DNA sample, 1 μL of each primer (2 µM), 5 μL of 2x KAPA HiFi HotStart ReadyMix (KAPA Biosystems, MA, USA), and 2 μL of ultrapure water. The PCR program was as follows: 95 °C for 3 min, 25 cycles at 95 °C for 30 sec, 55 °C for 30 sec, and 72 °C for 30 sec, and finally 72 °C for 5 min. The PCR products were purified using AMPure XP beads (Beckman Coulter, CA, USA). Each PCR product was indexed using the Nextera XT Index Kit (Nextera DNA UD Indexes set B; Illunima). The quality of libraries purified with AMPure XP beads was checked using a TapeStation D1000 (Agilent, CA, USA). Pooled libraries were sequenced using the Illumina MiSeq platform (Illumina), and 250 bp paired-end reads were generated (Table S1). Illumina raw reads were deposited in the NCBI SRA database under the accession number PRJXX0000000. Raw paired-end reads were analyzed using QIIME 2 (version 2020.8) (Bolyen et al., 2019) with the plugin “dada2” (Callahan et al., 2016) for quality filtering, trimming length, merging of paired reads, and removing chimeric sequences. Dada2-derived amplicon sequence variants (ASVs) with < 100 reads were excluded. Amplicon sequence variants (ASVs) with high sequence identity (> 99%) were manually combined. The resultant six ASVs were manually assigned to genus-level taxa using the BLAST function in the NCBI database.

### Fluorescence *in situ* hybridization (FISH) on aphid bacteriome and embryos

To visualize the localization and vertical transmission of *Wolbachia* in *C. cedri*, we conducted FISH on bacteriome and embryos with probes that were complementary to the 16S rRNA gene sequences (W1; 5’-Cy3-AATCCGGCCGARCCGACCC-3’ and W2; 5’-Cy3-CTTCTGTGAGTACCGTCATTATC-3’ [Heddi et al., 1999]). The two probes were used simultaneously to increase the signal. These probes were first developed for *Wolbachia* in *Sitophilus* weevils (Heddi et al., 1999) but were later confirmed to work well in *C. cedri* (Gómez-Valero et al., 2004). Aphids were collected three times from the campus of the National Institute for Basic Biology (NIBB) (Table 1). Fresh insects were immediately dissected in phosphate-buffered saline (PBS; 33 mM KH_2_PO_4_, 33 mM Na_2_HPO_4_, pH 7.4) under a stereomicroscope (SZ61; Olympus, Japan) with fine forceps. We dissected the ovarioles from late-instar nymphs or young adults of viviparous aphids to obtain a series of developing embryos. To collect bacteriomes, we dissected adults that possessed well-developed bacteriomes. Then collected ovaries and bacteriomes were fixed in 4% PFA in PBS for approximately 3 h. The fixed samples were washed thrice with PBS-Tx (0.3% Triton X-100 in PBS). Samples were also washed with hybridization buffer (20 mM Tris-HCl pH 8.0, 0.9 M NaCl, 0.01% SDS, and 30% [v/v] formamide) prior to hybridization. Then samples were incubated overnight at room temperature (25–28 □) in hybridization buffer containing the specific probes at a final concentration of 100 nM. During overnight incubation, DNA and F-actin were stained with 4,6-diamidino-2-phenylindole (DAPI) (Dojindo, Japan) and Alexa Fluor 488 phalloidin (Thermo Fisher Scientific, MA, USA), respectively. After hybridization, the samples were washed three times in PBS-Tx, mounted with VECTASHIELD (Vector Laboratories, CA, USA), and observed under a confocal laser scanning microscope FV1000 (Olympus). The acquired images were processed using image analysis software ImageJ (NIH, http://rsb.info.nih.gov/ij/). These observations were repeated three times.

## Results

### Microbiome and relative abundance of symbionts in *C. cedri*

To quantify the bacterial diversity associated with *C. cedri*, 16 individuals collected in Japan were subjected to 16S rRNA amplicon sequencing (Fig. 1, Table S1). *Wolbachia* were detected across all the populations, as well as *Buchnera* and *Serratia*, which are “obligate symbionts” in aphids (Fig. 1), which was consistent with a previous report on the USA populations (Jousselin et al., 2016). The relative abundance of *Buchnera, Serratia*, and *Wolbachia* was 51.19 ± 20.14, 39.24 ± 19.42, and 8.51 ± 4.15 (n = 16, mean % ± SD), respectively. The remaining minor bacterial taxa were not consistently detected and were revealed to be *Rosenbergiella, Phaseolibacter*, and *Klebsiella* allied species (Table S1).

**Fig. 1.**
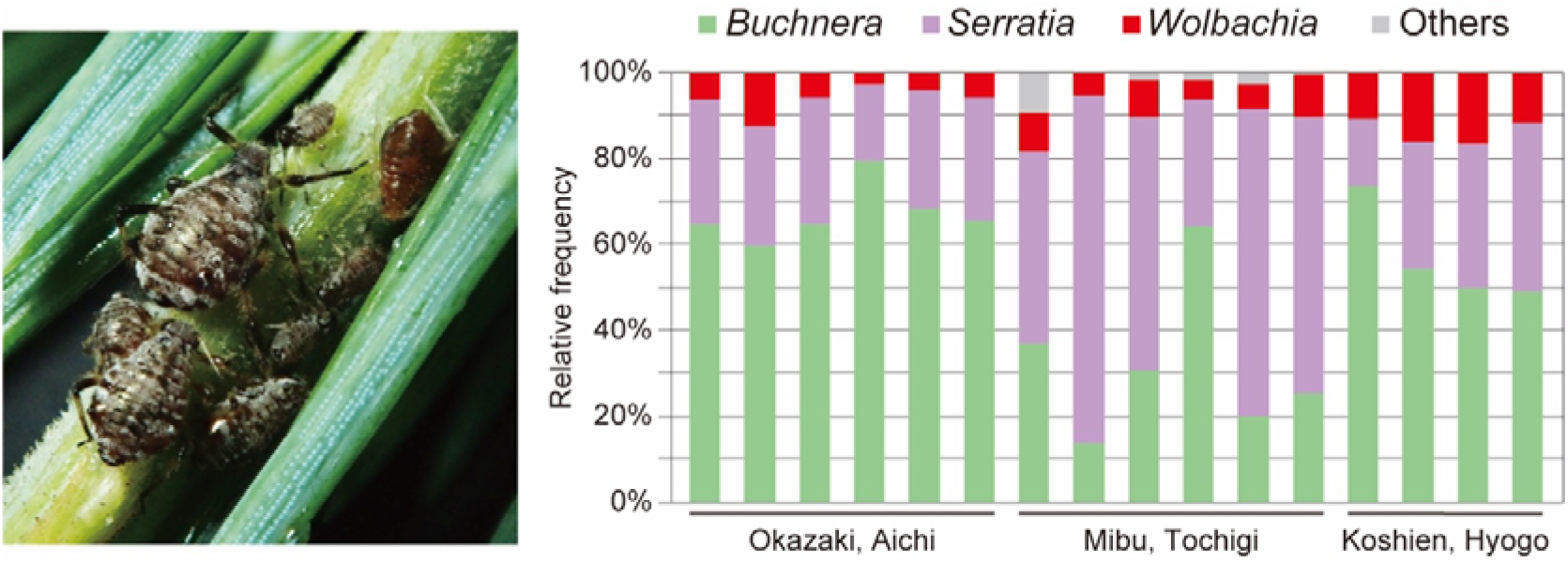
Left; image of *Cinara cedri* colony on twig of Himalayan cedar *Cedrus deodara*. Right; diversity in bacteria associated with *C. cedri*. Proportion of each bacterial species was quantified based on amplicon sequencing of hypervariable V3/V4 region of 16S rRNA gene. Assigned bacterial taxa (genus level) are color-coded as shown.

### Localization of *Wolbachia* endosymbiont in *C. cedri* bacteriome and embryo

*C. cedri* harbored three types of symbionts: *Buchnera, Serratia*, and *Wolbachia* (Fig. 1). To determine the localization of each symbiont, we conducted FISH with specific probes targeting 16S rRNA sequences. Based on our observations of the bacteriome in adults and late instar nymphs, we determined that *Buchnera* and *Serratia* are localized in distinct bacteriocyte types: *Buchnera* and *Serratia* bacteriocytes (Fig. S1). In contrast, *Wolbachia* signals were scattered throughout the cytoplasm of both the bacteriocytes and sheath cells (Fig. 2A). No signal was detected in intestinal or fat cells. This finding is consistent with a previous report (Gómez-Valero et al., 2004). We also observed *Wolbachia* localization in developing embryos. *Wolbachia* cells were recognized as clusters in the bacteriome region of abdominal part of the embryos (Fig. 2B).

**Fig. 2.**
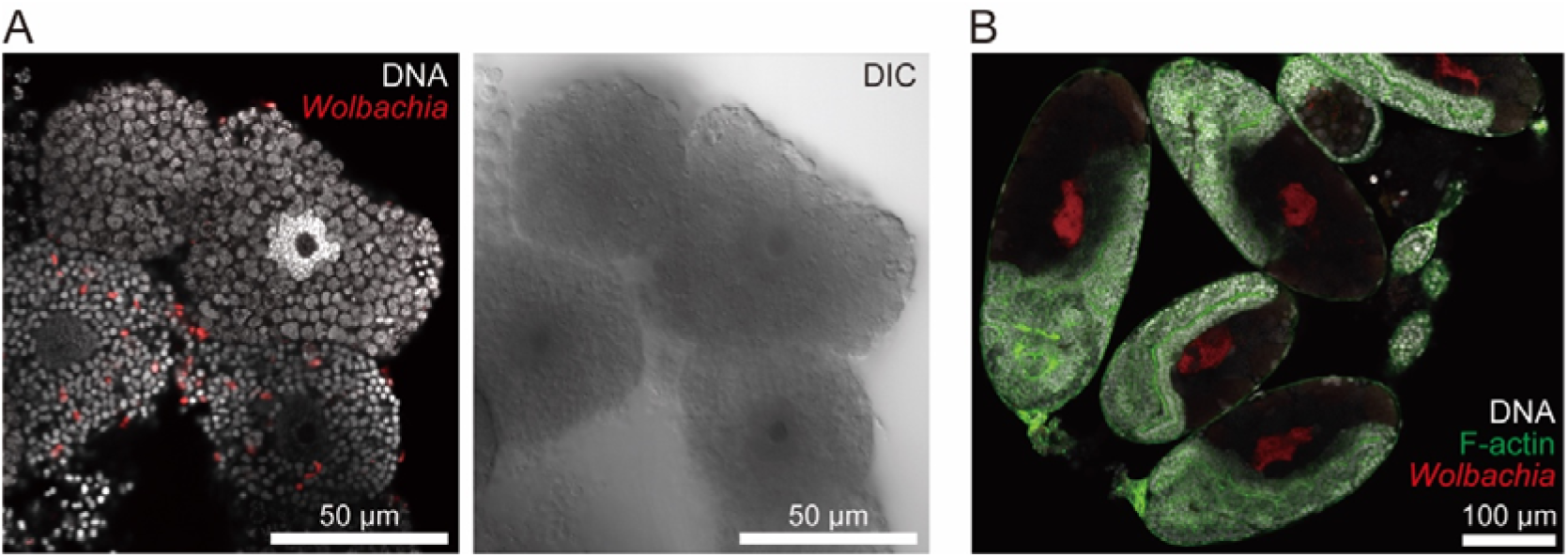
Localization of symbiotic bacteria in *Cinara cedri. Wolbachia* was visualized using FISH (red). **A** *Wolbachia* signals (red) were detected within bacteriocytes and sheath cells. *Wolbachia* cells were scattered throughout cytoplasm. DNA was stained with DAPI (white). **B** Signals were also detected in embryos. *Wolbachia* was recognized as a cluster within embryonic bacteriome. DNA and F-actin were stained with DAPI (white) and phalloidin (green), respectively.

### *Wolbachia* vertical transmission through viviparous ovarioles in *C. cedri*

To reveal the vertical transmission process of *Wolbachia*, we conducted FISH observations at various stages of embryo development (for developmental categories, see Fig. S4, S5, and SI text). We categorized each embryo of *C. cedri* into eight developmental stages: oocyte, syncytial blastoderm, cellular blastoderm stages I and II, invagination, segmentation, flip, and final growth (Fig. S2, S3). Based on these categories, we analyzed the symbiont infection stages and described the infection process of *Wolbachia*.

At the oocyte stage, when the oocytes were separated from the germarium, *Wolbachia* signals were observed in the interstitial spaces of nurse cells; however, this was not consistent (germaria without *Wolbachia* signals were frequently observed) (Fig. 3A). In the syncytial blastoderm and early cellular blastoderm stages, there was no signal of *Wolbachia* (Fig. 3B). *Wolbachia* signals were consistently detected in the later blastoderm stage, in which both *Buchnera* and *Serratia* obligate symbionts began to be transmitted from the mother’s bacteriocytes (Fig. 3C). All three symbionts clustered, and the mixed population was incorporated into the embryo from the posterior region. Symbiont incorporation continued through subsequent invagination (anatrepsis) and segmentation stages. After the uptake of the symbionts, *Buchnera* and *Serratia* were enclosed as bacteriocytes and began to proliferate, whereas *Wolbachia* cells were scattered within the bacteriocytes or clustered in the spaces between them (Fig. 4D).

**Fig. 3.**
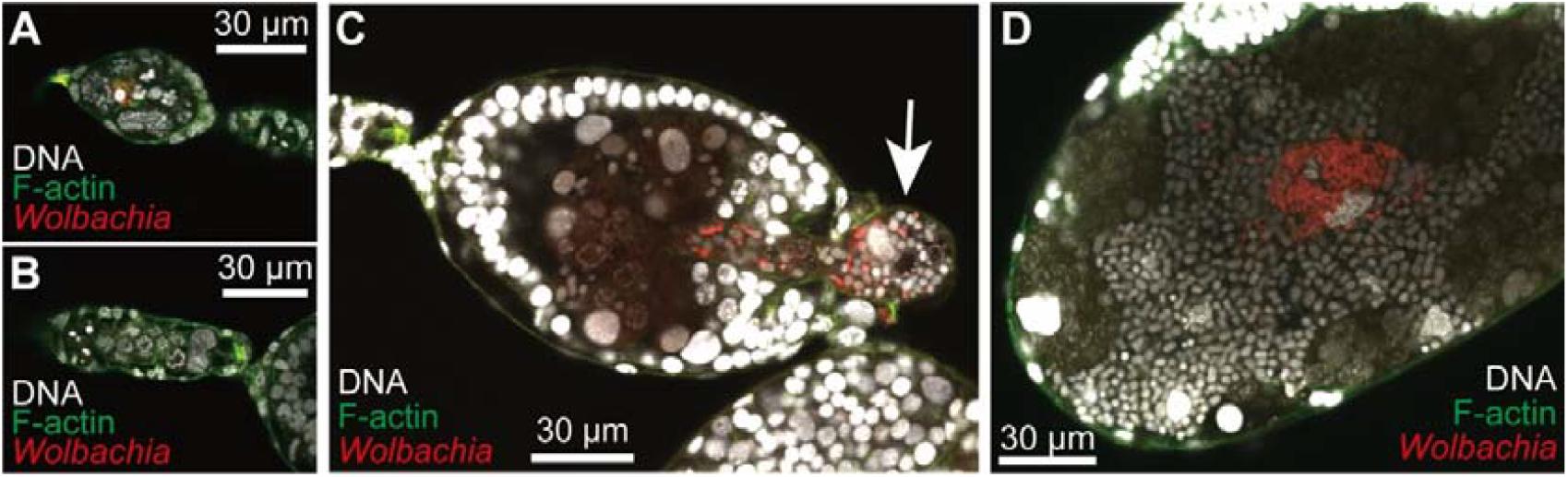
*Wolbachia* vertical transmission in *Cinara cedri. Wolbachia* were visualized using FISH (red), and DNA and F-actin were stained with DAPI (white) and phalloidin (green), respectively. **A** *Wolbachia* invaded into germ cells but their infection rates were not high. **B** *Wolbachia* cells were rarely observed in embryos at early stages, prior to transmission of main symbiotic bacteria (*Buchnera* and *Serratia*). **C** *Wolbachia*, along with obligate symbionts, were simultaneously transmitted from maternal tissues to embryo. White arrow indicates mass of symbionts at “entry point”. **D** After transmission, number of *Wolbachia* cells were noticeably increased in limited area of embryonic bacteriome.

## Discussion

A hallmark of *Wolbachia* is its efficient and stable maternal transmission coupled with its ability to induce diverse host phenotypes, contributing to its global prevalence (Kaur et al., 2021; Pietri et al., 2016). The transmission route and its underlying mechanism are well-studied in some *Wolbachia* infecting oviparous insects; *Wolbachia* first localize in the host maternal germline and then, transmitted to oocytes, through the connection between nurse cells and early stages of oocytes, and/or by cooption of the uptake machinery of yolk proteins (Guo et al., 2018; Russel et al., 2019; Serbus & Sullivan 2007). However, in the present study, we demonstrated that the *Wolbachia* endosymbiont in the aphid *C. cedri* was directly transmitted from the maternal bacteriome to developing embryos, together with the obligate symbionts *Buchenra* and *Serratia* (Fig. 3). Consistent with a previous report, *Wolbachia* in *C. cedri* was consistently detected (Fig. 1; Augustinos et al., 2011; Jousselin et al., 2016; Nozaki et al., 2022) and localized in the maternal bacteriome and embryos (Fig. 2; Gómez-Valero et al., 2004). Although *Wolbachia* signals were detected in some germaria, their absence in the earliest stage of embryogenesis indicated that transfer from germaria to oocytes did not occur (Fig. 3). These results suggest that *Wolbachia* in *C. cedri* piggybacked on the transmission route is commonly utilized by aphid obligate symbionts and does not adhere to the typical strategy of this bacterial genus. Further observation of the transmission of *Wolbachia* in other aphid species or different strains of *Wolbachia* in *C. cedri* can clarify whether this modification of the transmission pathway is caused by the reproductive specificities of the host insect or by bacterial factors.

*Wolbachia* in *C. cedri* exhibited direct transmission from maternal cells in the bacteriome to offspring with obligate symbionts (Fig. 3), which is not typical of the bacterium. This deviation may be attributed to the rapid viviparous oogenesis in aphids, which is characterized by a shortened vitellogenic growth phase. Viviparous oocytes are much smaller than sexual aphids, and the connection between germarium nurse cells and oocytes is tightly restricted (Bickel et al., 2013; Miura et al., 2003). Unlike oviparous insects, there may be simply not enough time for *Wolbachia* to infect oocytes from nurse cells in the germarium. However, our additional observations of the oviparous ovarioles in this aphid contradicted this hypothesis; *Wolbachia* were not localized in germarium cells and were transmitted together with obligate symbionts (SI text and Fig. S6). Therefore, factors other than viviparous oogenesis, such as *Wolbachia*’s ability to target and interact with diverse cellular components and organelles (Porter and Sullivan, 2023), or aphid immune responses (Zug and Hammerstein, 2015), warrant future genomic and experimental investigations.

In the present study, we exploited an introduced population of *C. cedri* in Japan (Nozaki et al., 2022), and successfully observed the transmission of symbionts in detail. In Japan, only *Wolbachia* supergroup M, an aphid-specific clade, has been detected (Nozaki et al., 2022), whereas supergroup B has also been reported in the eastern Mediterranean, a potential native range (Augustinos et al., 2011; Michelena et al., 2005). Investigating the vertical transmission routes of other *Wolbachia* strains in this aphid species will clarify whether the observed modifications in *Wolbachia* transmission are bacterial lineage-specific. It is also worth elucidating the transmission of the M-supergroup strain of *Wolbachia* in banana aphids, which is the best-studied aphid group in terms of its association with *Wolbachia* (De Clerck et al., 2014; Higashi et al., 2024; Jones et al., 2011).

## Conclusion

In summary, we report a novel *Wolbachia* transmission strategy in *C. cedri*, where *Wolbachia* utilizes the obligate symbiont transmission pathway, deviating from the canonical germline-to-oocyte transfer. Future genomic and experimental studies, including comparative genomics among related *Wolbachia* species and artificial infection experiments, are crucial for elucidating the mechanisms driving this unique adaptation. These findings underscore the diversity of *Wolbachia* transmission strategies and provide a framework for future studies on the molecular determinants of endosymbiont vertical transmission, including targeting/migration capabilities of *Wolbachia* (Porter and Sullivan, 2023), and interactions with other symbionts and the host (Bright and Bulgheresi, 2010; Vautrin and Vavre, 2009; Zug and Hammerstein, 2015).

## Data Availability Statement

Raw reads (FASTQ files) were deposited in the SRA (NCBI). The BioProject accession number is PRJXX0000000.

## Acknowledgements

We thank Dr. Shunta Yorimoto for sample collection, and Mika Ikeda for technical support. We also appreciate the members of Laboratory of Evolutionary Genomics, National Institute for Basic Biology (NIBB) and the Functional Genomics Facility, NIBB Core Research Facilities. Computational resources were provided by the Data Integration and Analysis Facility, NIBB.

## Conflicts of Interest

The authors declare no conflicts of interest.

## Author Contributions

T.N. was involved in conceptualization, data curation, formal analysis, funding acquisition, investigation, methodology, project administration, resources, validation, visualization, writing—original draft, and writing—review and editing. Y.K. was involved in data curation, investigation, methodology, resources, writing—original draft, and writing—review and editing. S.S. was involved in the conceptualization, funding acquisition, investigation, project administration, writing—original draft, and writing—review and editing. All authors provided their final approval for publication and agreed to be held accountable for the work performed.

## Funding

This work was financially supported by the Japan Society for the Promotion of Science awarded to T.N. (Research Fellowship for Young Scientists Nos. 19J01756 and KAKENHI 22K14901) and S.S. (KAKENHI 17H03717, 17H06384 and 20H00478).

## Ethics statement

The insect samples used in this study were collected from the wild. The experimental methods employed do not require ethical approval.

## Notes

### Competing Interest Statement

The authors have declared no competing interest.

