## Supplementary material for "Vertical transmission of *Wolbachia* bypassing the germ line in an aphid": SI

**This file contains:**
**SI Text (Methods, and Result and Discussion)**
**Table S1**
**Figure S1–S4**

### SI Text

#### SI Methods

##### Structure of *Cinara cedri* bacteriome cells

Generally, aphid bacteriomes consist of two types of cells: bacteriocytes containing the obligate symbiont such as *Buchnera* in their cytoplasm and sheath cells without them (Nozaki & Shigenobu, 2022). Bacteriocytes are large polyploid cells (~256 ploidy in *Acyrthosiphone pisum*) that are recognized as the interface for nutritional interactions with symbiotic bacteria. In some aphids with "multiple obligate symbionts," such as *Cinara cedri* and *Ceratovacuna japonica*, each obligate symbiotic bacterium occupies distinct bacteriocytes (Gómez-Valero et al., 2004; Yorimoto et al., 2022). Sheath cells often harbor facultative symbionts (Moran et al., 2005; Koga et al., 2012; Yorimoto et al., 2022). To gain more detailed information on the cellular features of *C. cedri* bacteriome, we performed a morphological analysis of the dissected bacteriomes.

To visualize the obligate symbionts, we conducted fluorescent *in situ* hybridization (FISH) on the bacteriome of *C. cedri*, using symbiont-specific probes targeting the 16S rRNA gene sequences (*Buchnera aphidicola* Cc: 5'-FAM-CCCGTTCGCCGCTCGCCGGCA-3' [Gosalbes et al., 2008], *Serratia symbiotica* Cc: 5'-Cy3-CCGCCGCTCGTCACCCAAA-3' [Manzano-Marin et al., 2016]). Viviparous individuals collected from the campus of the National Institute for Basic Biology (NIBB) were immediately dissected in phosphate-buffered saline (PBS; 33 mM $KH_2PO_4$ and 33 mM $Na_2HPO_4$, pH 6.8) under a stereomicroscope (SZ61; Olympus, Japan) with fine forceps. Bacteriomes were dissected from young adults and fixed in 4% paraformaldehyde in PBS for approximately 3 h. The fixed samples were washed three times with PBS-Tx (0.3% Triton X-100 in PBS), and then with hybridization buffer (20 mM Tris-HCl pH 8.0, 0.9 M NaCl, and 0.01% SDS, 30% [v/v] formamide) before hybridization. The samples were incubated overnight at room temperature (25–28 ℃) in hybridization buffer containing two specific probes at a final concentration of 100 nM and 1 μg/mL of 4,6-diamidino-2-phenylindole (DAPI) (Dojindo, Japan) for DNA staining. After overnight incubation, the samples were washed thrice with PBS-Tx, mounted with VECTASHIELD (Vector Laboratories, CA, USA), and observed under a confocal laser scanning microscope FV1000 (Olympus). To determine the structure of the bacteriomes, we also stained, dissected, fixed, and washed them with DAPI (1 μg/mL; Dojindo) for the nuclei and Alexa Fluor™ 488 phalloidin (66 nM; Thermo Fisher Scientific, MA, USA) for the cytoskeleton (F-actin), respectively.

##### Embryonic development and transmission of obligate symbionts in *C. cedri*

To determine the vertical transmission of the *C. cedri* obligate symbionts *Buchnera* and *Serratia*, we observed viviparous ovarioles harboring a series of developmental embryos and bacteriomes using FISH. The experimental procedures were performed as described above. The morphology of the embryos and symbiont localization were recorded. We also measured the lengths of the major and minor axes of the embryos to estimate their volume using ImageJ software (NIH, http://rsb. info. nih. gov/ ij/). Through observations, we described the embryonic development of this aphid and determined the entry site/time and behavior of both symbionts, with reference to detailed information from the well-studied aphid *A. pisum* (Miura et al., 2003).

**Observation on *Wolbachaia* in oviparous ovarioles of *C. cedri***

In January 2022, two adult individuals with both oviparous and viviparous ovarioles (possibly morph-mosaicism) were collected from the NIBB campus. Although they were abnormal individuals, we took this finding as a good opportunity to observe their vertical transmission through the oviparous ovarioles. Ovarioles with oocytes were processed and hybridized with *Wolbachia* probes as described in the main text.

#### SI Results and Discussion

##### Staging aphid viviparous development

In this study, we present the first description of viviparous embryonic development with the transmission of endosymbiotic bacteria in *C. cedri* (Fig. S2). Based on the observations of 117 embryos, we characterized eight developmental stages: 1) oocyte stage; 2) syncytial blastoderm stage; 3) cellular blastoderm stage I; 4) cellular blastoderm stage II; 5) invagination stage; 6) segmentation stage; 7) flip stage; and 8) final growth stage (Fig. S2). Technical terms and general descriptions followed those described by Miura et al. (2003). The increase in embryo size at the embryonic developmental stage suggests that this stage division is generally reasonable (Fig. S3).

Embryonic development and symbiont transmission have been well studied in the pea-aphid *A. pisum* (Miura et al., 2003; Braendle et al., 2003; Koga et al., 2012). The embryogenesis of *C. cedri* broadly aligned with that of *A. pisum* (Fig. S2) while detailed and direct stage-by-stage comparisons were not feasible. However, the time window for *C. cedri* embryos to become receptive to symbiont infections was longer than that for *A. pisum*. In *A. pisum*, transmission ends before invagination of the germband (anatrepsis) begins (Miura et al., 2003), but in *C. cedri*, transmission continues even when the embryo becomes more elongated and segmented (from stages 4 to 6 in *C. cedri*; Fig. S2). This pattern is similar to the symbiont transmission in *Ceratovacuna japonica*, which harbors two obligate symbionts: *Buchnera* and *Arsenophonus* (Yorimoto et al., 2022). Notably, in both *C. cedri* and *Ceratovacuna japonica*, bacteriocyte formation and symbiont transmission overlapped; these bacteria were internalized within bacteriocytes (*Buchnera* and *Serratia* in distinct bacteriocytes) prior to transmission completion (Fig. S2; Yorimoto et al., 2022). In multi-partner symbiotic systems, there must be a precise mechanism for the segregation of multiple bacterial species into distinct host embryos/cells. Investigating this mechanism could provide novel insights into the microbial control systems within organisms.

##### ***Wolbachia* vertical transmission through oviparous ovarioles in *C. cedri***

We had unique opportunity to observe oviparous ovarioles in the morph-mosaic females. Notably, *Wolbachia* cells were absent from the germarium, suggesting a transition from the maternal bacteriome to the developing oocyte along with *Buchnera* and *Serratia* (Fig. S4). This was consistent with the observation of viviparous development (Fig. 3). However, we must emphasize that further studies are necessary to validate our observations. Specifically, we cannot exclude the possibility that the atypical developmental patterns observed during winter were a consequence of a recent introduction to Japan, where environmental conditions may have diverged significantly from the native habitat.

#### SI Table

**Table S1.** Detailed information of 16S rRNA amplicon sequencing.

| # | Insect stage (vivi. female) | Locality | No. of raw read* | BioSample No. (PRJXX0000000) | Detected bacteria[†] |
|---|---|---|---|---|---|
| 1 | Adult | Okazaki, Aichi | 192,277 | XX0000000 | ***Buchnera***, ***Serratia***, ***Wolbachia*** |
| 2 | Adult | Okazaki, Aichi | 180,832 | XX00000007 | ***Buchnera***, ***Serratia***, ***Wolbachia*** |
| 3 | Adult | Okazaki, Aichi | 146,704 | XX0000000 | ***Buchnera***, ***Serratia***, ***Wolbachia*** |
| 4 | Adult | Okazaki, Aichi | 212,665 | XX0000000 | ***Buchnera***, ***Serratia***, ***Wolbachia*** |
| 5 | Adult | Okazaki, Aichi | 212,254 | XX0000000 | ***Buchnera***, ***Serratia***, ***Wolbachia*** |
| 6 | Adult | Okazaki, Aichi | 200,090 | XX0000000 | ***Buchnera***, ***Serratia***, ***Wolbachia*** |
| 7 | Adult or late-stage nymph | Mibu, Tochigi | 21,188 | XX0000000 | ***Buchnera***, ***Serratia***, ***Wolbachia***, ***Rosenbergiella***, *Klebsiella* |
| 8 | Adult or late-stage nymph | Mibu, Tochigi | 24,146 | XX0000000 | ***Buchnera***, ***Serratia***, ***Wolbachia***, *Rosenbergiella* |
| 9 | Adult or late-stage nymph | Mibu, Tochigi | 22,538 | XX0000000 | ***Buchnera***, ***Serratia***, ***Wolbachia***, ***Rosenbergiella***, *Phaseolibacter* |
| 10 | Adult or late-stage nymph | Mibu, Tochigi | 29,335 | XX0000000 | ***Buchnera***, ***Serratia***, ***Wolbachia***, ***Rosenbergiella***, *Phaseolibacter* |
| 11 | Adult or late-stage nymph | Mibu, Tochigi | 23,377 | XX0000000 | ***Buchnera***, ***Serratia***, ***Wolbachia***, ***Rosenbergiella***, *Phaseolibacter* |
| 12 | Adult or late-stage nymph | Mibu, Tochigi | 26,284 | XX0000000 | ***Buchnera***, ***Serratia***, ***Wolbachia***, *Rosenbergiella* |
| 13 | Late-stage nymph | Nishinomiya, Hyogo | 17,058 | XX0000000 | ***Buchnera***, ***Serratia***, ***Wolbachia*** |
| 14 | Late-stage nymph | Nishinomiya, Hyogo | 16,777 | XX0000000 | ***Buchnera***, ***Serratia***, ***Wolbachia*** |
| 15 | Late-stage nymph | Nishinomiya, Hyogo | 19,907 | XX0000000 | ***Buchnera***, ***Serratia***, ***Wolbachia*** |
| 16 | Late-stage nymph | Nishinomiya, Hyogo | 17,711 | XX0000000 | ***Buchnera***, ***Serratia***, ***Wolbachia*** |

* 250 bp of paired-end reads generated by Illumina MiSeq.

[†] Bacterial taxa detected with 100 or more reads as ASV in at least one sample are listed. Those accounting for more than 1% of the total reads per sample are shown in **bold**.

#### SI Figures

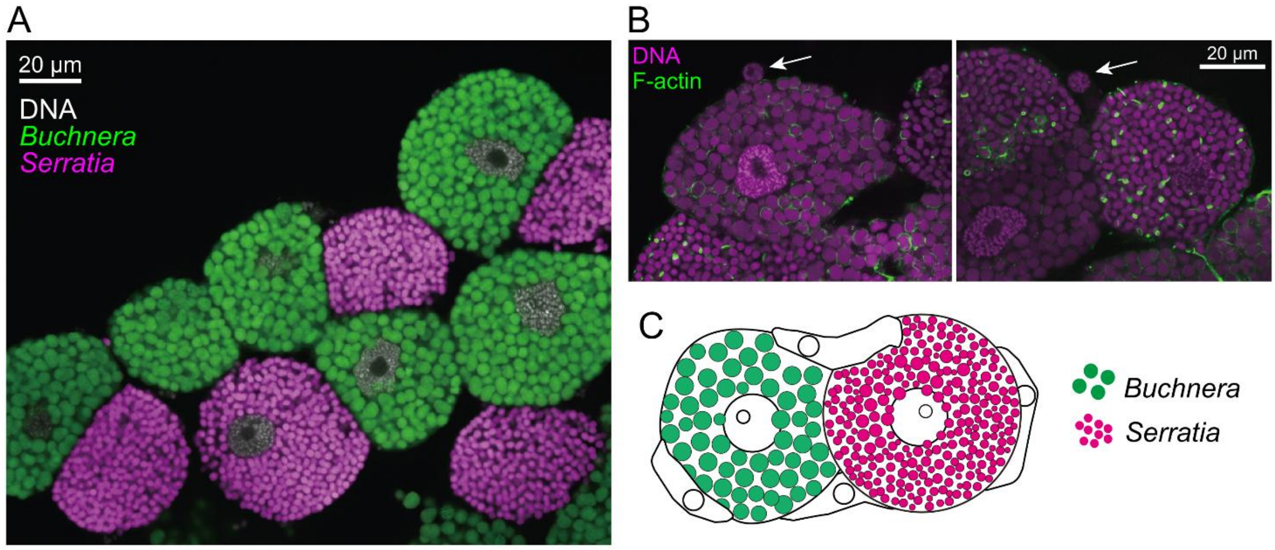


**Fig. S1**. Localization of obligate symbionts and structure of bacteriome in *Cinara cedri*. **A** Localization of *Buchnera* (green) and *Serratia* (magenta) visualized with fluorescent *in situ* hybridization targeting of 16S rRNA of each symbiont. Bacteriocytes containing *Buchenra* and *Serratia* were distributed in a nested manner. **B** Morphology of bacteriome visualized with DAPI-phalloidin staining. DNA and F-actin were stained with DAPI (magenta) and phalloidin (green), respectively. Sheath cells (arrows) were observed between bacteriocytes and on surface. **C** Schematic illustration of *C. cedri* bacteriome, which is consisted with two types of bacteriocytes (*Buchnera* and *Serratia*) and sheath cells.

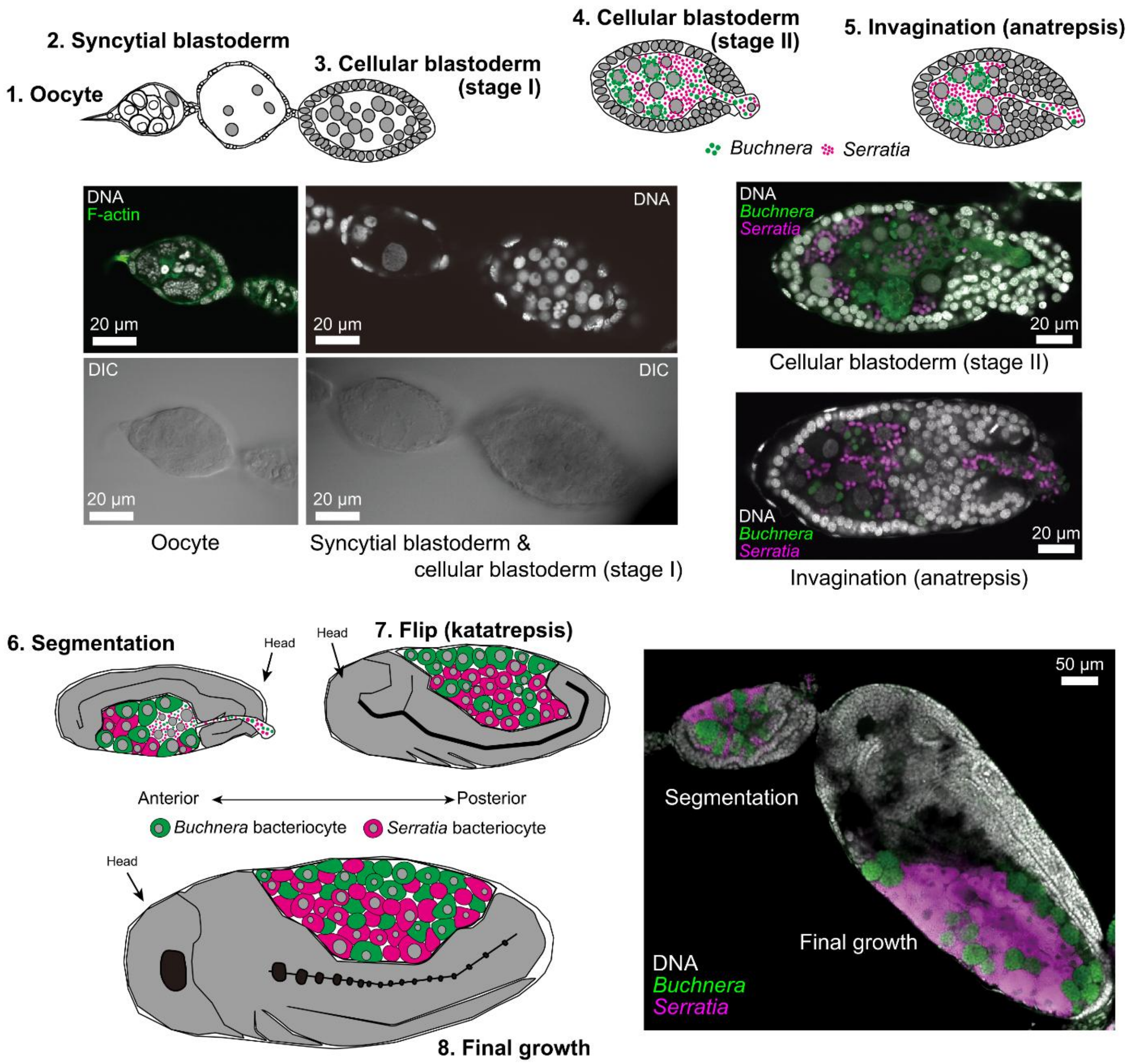


**Fig. S2**. Vertical transmission of obligate symbionts and bacteriome formation in *Cinara cedri*. Embryonic development of *C. cedri* was divided into eight stages: oocyte (1), syncytial blastoderm (2), cellular blastoderm stages I (3) and II (4), invagination (5), segmentation (6), flip (7), and final growth (8). Embryogenesis of *C. cedri* was generally similar to that of *Acyrthosiphone pisum* (Miura et al., 2003) and *Ceratovacuna Japonica* (Yorimoto et al., 2022), although the stages we described in the current study cannot be simply compared with those in other species.

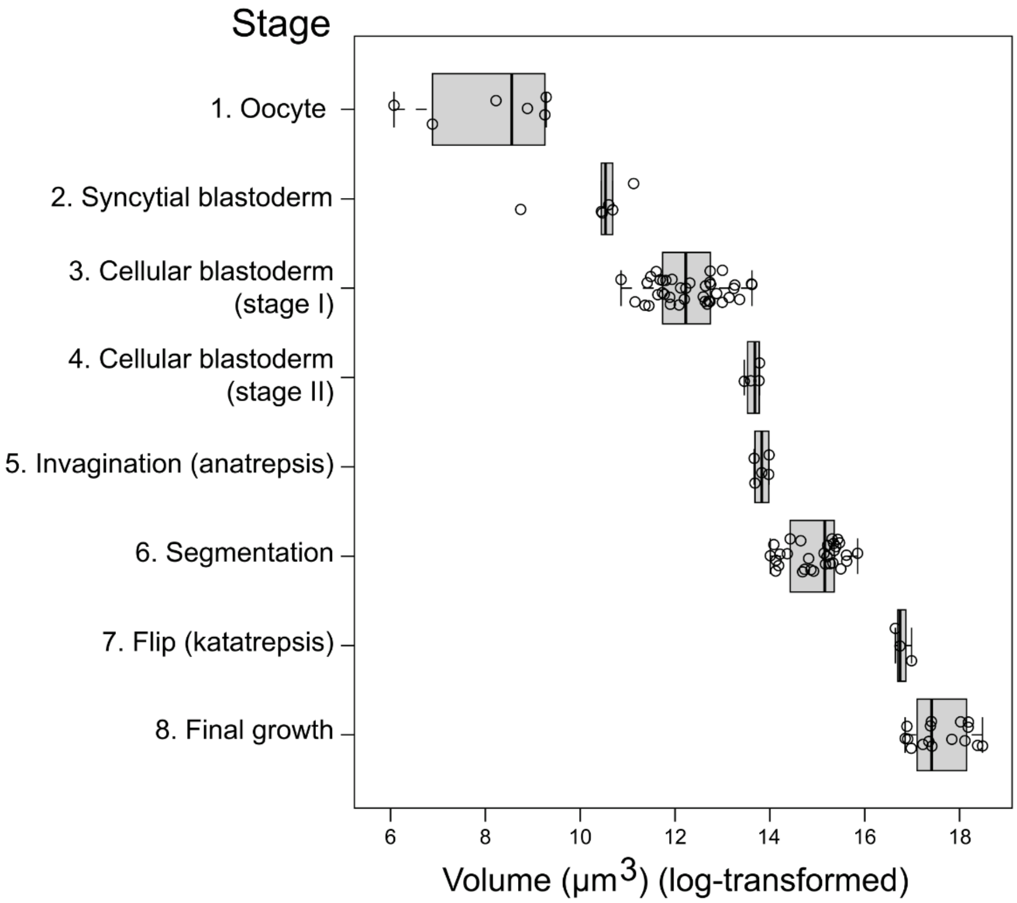


**Fig. S3.** Embryo size change associated with developmental stages. Approximate egg volumes were calculated using the formula: $V = 4\pi\left(\frac{L}{2}\right)\left(\frac{W}{2}\right)\frac{2}{3}$ , where $L$ and $W$ are long and short axes, respectively. Embryos were categorized into eight stages based on our observations (Fig. S2).

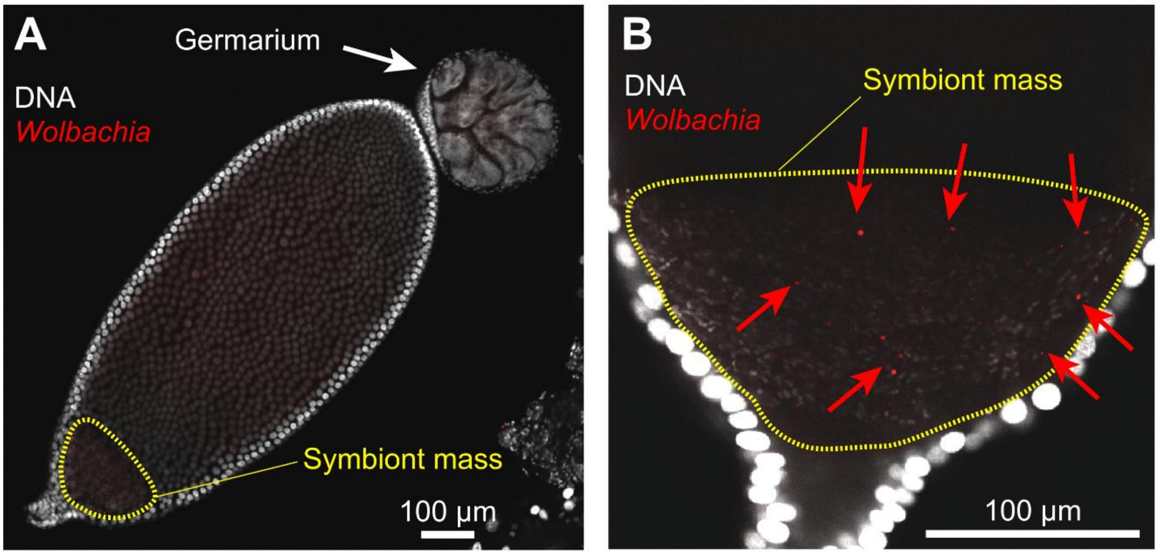


**Fig. S4.** Fluorescence *in situ* hybridization (FISH) of *Wolbachia* in oviparous ovarioles of *Cinara cedri.* Oviparous ovarioles from two morph-mosaic females (exhibiting both viviparous and oviparous morphs) were examined using FISH to visualize *Wolbachia* localization. **A** An oviparous ovariole showing germarium (cluster of germ and nurse cells) at anterior tip, where *Wolbachia* signal was absent. Developing oocytes displayed a symbiont mass (demarcated by a yellow dashed line) in the posterior region. **B** Magnified view of symbiont mass region, revealing *Wolbachia* cells (red dots, indicated by red arrows) interspersed with DAPI-positive particles (reasonably assuming *Buchnera* and *Serratia*).
